# High throughput proteomics identifies 484 high-accuracy plasma protein biomarker signatures for ovarian cancer

**DOI:** 10.1101/349829

**Authors:** Stefan Enroth, Malin Berggrund, Maria Lycke, John Broberg, Martin Lundberg, Erika Assarsson, Matts Olovsson, Karin Stålberg, Karin Sundfeldt, Ulf Gyllensten

**Affiliations:** Department of Immunology, Genetics, and Pathology, Biomedical Center, Science for Life Laboratory (SciLifeLab) Uppsala, Box 815, Uppsala University, SE-75108 Uppsala, Sweden; Department of Obstetrics and Gynaecology, Institute of Clinical Sciences, Sahlgrenska Academy at Gothenburg University, Gothenburg, Sweden; OLINK Proteomics, Uppsala Science Park, SE-751 83, Uppsala, Sweden; Department of Women’s and Children’s Health, Uppsala University, Uppsala, Sweden

**Keywords:** Biomarker, Ovarian cancer, benign tumours, plasma protein

## Abstract

Ovarian cancer is usually detected at a late stage with the 5-year survival at only 30-40%. Additional means for early detection and improved diagnosis are acutely needed. To search for novel biomarkers, we compared circulating plasma levels of 981 proteins in patients with ovarian cancer and benign tumours, using the proximity extension assay. A novel combinatorial strategy was developed for identification of multivariate biomarker signatures, resulting in 484 mutually exclusive models out of which 448 did not contain the present biomarker MUCIN-16. The top-ranking model consisted of 14 proteins and had a AUC=0.95, PPV=1.0, sensitivity=0.99 and specificity=1.0 for detection of stage III-IV ovarian cancer in the discovery data, and an AUC=0.89, PPV=0.93, sensitivity=0.89 and specificity=0.95 in the replication data. The novel plasma protein signature could be used to improve the diagnosis of women with adnexal ovarian mass or in screening to identify women that should be referred to specialized examination.

## Introduction

Ovarian cancer is currently the 7^th^ most common cancer across the world with estimated incidences from 4.1 to 11.4 cases per 100 000 women^1^. Since ovarian cancer is commonly caught late, the overall 5-year survival rate is only 30-40%. MUCIN-16 (also known as Cancer antigen 125, CA-125) was introduced as a biomarker for ovarian cancer in 1983^2^ and is currently the most important single biomarker for epithelial ovarian cancer managment^3^. MUCIN-16 alone however, has low sensitivity for early stage cancer (50-62%) at a specificity of 94-98.5%^3^. Combinations of MUCIN-16 and other biomarkers, including WFDC2 (WAP Four-Disulfide Core Domain 2, also known as HE4 - human epididymal protein 4), such as the ROMA Score (Ovarian Malignancy Risk Algorithm), increases the sensitivity to 75% at similar specificity (90-95%)^4^. The low sensitivity for detection of early stage ovarian cancer prohibits population screening using the current biomarkers. A recent study in the UK suggests that multi-modal tests are approaching sufficient accuracy to justify screening from a health-economic stand-point^5^. However, tests with low specificity have a high false positive rate, which will result in unnecessary anxiety and examinations and also additional cost for the health-care system.

The presently available biomarkers are mainly used to improve diagnosis of women that experience symptoms or when imaging such as transvaginal ultrasound (TVU) or computer tomography (CT) indicate adnexal ovarian mass. The tests/algorithms then triage patients in need of surgery at tertiary centres. Even in this context, identification of clinically useful biomarkers based on single or combination of proteins is challenging. Recent developments of high-throughput technologies for detection and quantification of proteins has made it possible to study thousands of biomarker candidates in a single sample. Skates and colleagues^6^ have presented a statistical framework for study design, sample size calculation in discovery and replication stages, for identification of single biomarkers that can distinguish between cases and controls, with special reference to ovarian cancer. They recommend selection of the highest ranking 50 biomarkers from a discovery stage, which are then examined in a replication stage. A smaller set of replicated markers is then used to build a classifier that is tested in clinical validation studies. We have previously shown^7^ that plasma protein levels for several protein biomarkers are highly correlated. This implies that sets of proteins can be identified in a discovery stage whose combined predictive power is not greater than their individual contribution. Also, biomarkers that are not significant on their own can increase the predictive power in combination with other, individually significant or non-significant, biomarkers.

One approach for finding combinations of highly predictive biomarkers is to use exhaustive searches, such as the approach taken by Han and colleagues^8^ where 165 combinations of MUCIN-16 and a selection of three out of 11 additional biomarkers were examined for their ability to separate high-grade serous ovarian carcinoma from benign conditions. Such exhaustive approaches quickly become computationally unfeasible when the number of candidate proteins is high. For instance, choosing 4 from 1000 proteins can be done in over 40 billion ways. Another strategy is to use feature selection with machine learning frameworks to select subsets of informative markers from a larger set. This approach has previously been used to build a classifier with 12 biomarkers selected from 92 for separating ovarian cancer from healthy controls or benign conditions^9^. This is achieved by splitting the samples into a training and a test set, but with fairly small sample sizes different models are usually generated depending on the subset of samples used for training. To overcome these limitations, we developed a novel analysis strategy based on building models separating ovarian cancer from benign tumours, where we first identify smaller sets of proteins that are robustly selected across several splits, so-called cores. In the second step, we build a model by extending a core with additional proteins that have high predictive power in combination with the specific core.

Here, we aimed to identify multiple mutually exclusive biomarker signatures differentiating benign conditions from ovarian cancers at different stages, grades and all histological subtypes. The signatures should be practically useful and contain up to 20 proteins selected from a total of 981 characterized plasma proteins in one discovery cohort and two replication cohorts.

## Results

### Characterization of plasma proteins in the discovery and replication cohorts

A total of 552 proteins were characterized in the discovery and replication cohorts using the proximity extension assays (PEA) with the Olink Proseek Cardiometabolic, Cell Regulation, Development, Immune Response, Metabolism and Organ Damage panels (Methods). These measurements were combined with data from a previous study [Enroth et al, unpublished] on 460 characterized proteins in the discovery cohort, bringing the total number of unique proteins included in the analysis to 981. Forty-two of the 460 proteins have also been quantified in the replication cohorts using the proximity extension assay in two custom 21-plex panels as previously described [Enroth et al, unpublished]. Following quality controls and normalization (Methods), a common set of 593 proteins (42 proteins from the previous 5 panels and 551 from the additional 6 panels) characterized in all samples were used.

### 484 distinct predictive models for ovarian cancer

Models were generated using only the discovery data, according to our two-stage strategy. First, mutually exclusive protein cores, consisting of a smaller set of proteins, were selected by repeatedly splitting the data into training and test sets and retaining proteins that were present in at least 70% of the models (Material and Methods, Figure 1A, B). Additional biomarkers where subsequently added to each core using a stepwise forward selection approach (Material and Methods, Figure 1C). The addition of proteins where terminated when the total model size was 20 proteins, or the next protein to be added did not substantially increase the performance of the model (Material and Methods). Using this strategy, we generated models to distinguish benign tumours from ovarian cancer stages I-II, III-IV and I-IV. For benign tumours versus stages I-II, the core had to be 2-6 proteins in length and have a sensitivity of at least 0.8, or a sensitivity and specificity above 0.6. For stages III-IV, the allowed core size was 2-5, and had to have a sensitivity above 0.8 or a sensitivity and specificity above 0.7. Finally, for stages I-IV the allowed core size was 2-6 proteins, and the models were required to have a sensitivity above 0.8 or a sensitivity and specificity above 0.7. These different parameter settings were used to identify models that can have either high sensitivity or high specificity or both, depending on the final application and to account for the fact that it is much more difficult to separate stages I and II from benign tumours compared to stages III and IV.

**Figure 1.**
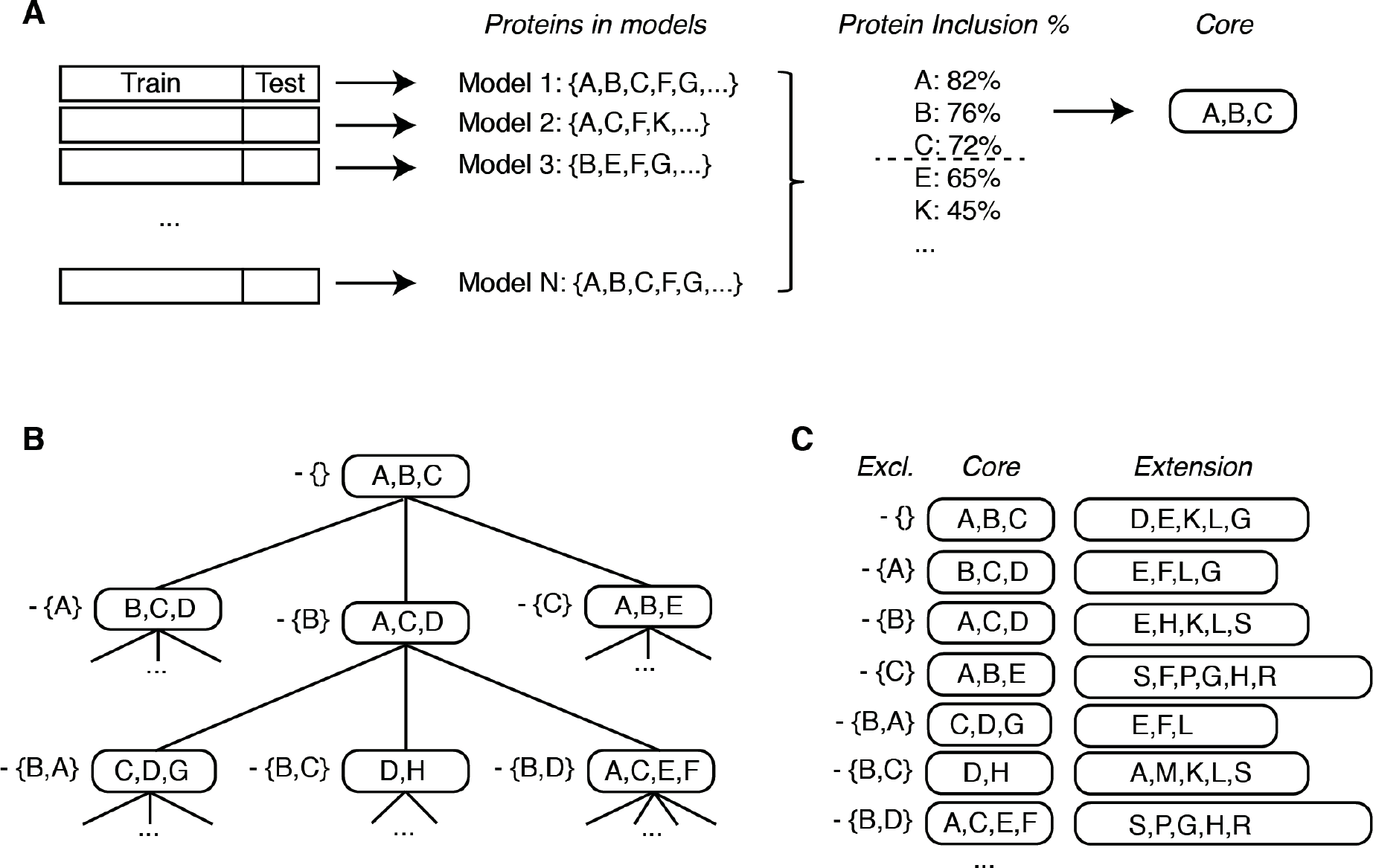
Model Generation. **(A)** Repeated model generation over random splits of the data. Proteins present in a sufficient fraction of the models are included into the core. **(B)** Generation of mutually exclusive cores. Proteins present in the first core (top node) are sequentially withheld from the second round of core discovery, as indicated by the sets to the left of the nodes. Each core of size N generates N new search-branches. **(C)** The final models are built by adding proteins to each core. The added proteins are chosen with respect to the proteins excluded in the core-discovery. Proteins are added in a stepwise forward selection choosing the protein that explains the highest proportion of remaining variance in the decision. See Materials and Methods for details.

The analysis resulted in 484 unique, mutually exclusive, models. The individual performance in the test-partition of the discovery data for the highest ranking 50 models is shown in Figure 2A. MUCIN-16, which is the clinically most useful single biomarker today, was the most common protein across cores in the 50 highest ranking models when sorting on core-performance (average sensitivity and specificity in the test set from the discovery data, Figure 2B). Our search strategy specifically excludes sets of protein, and 448 of the detected cores did not contain MUCIN-16. In general, when MUCIN-16 was not included, the models contained a higher number of proteins (9 to 20) than when it was included (8 to 17). Overall, 371 proteins were included in a core, or as an additional protein in at least one model. Among the top-ranking 50 cores and models, 115 proteins were selected in the addition phase compared to the list of 19 proteins that made up the core-set (Figure 2B, C). The performance of the 484 models in the test data is listed in Table 2 and a complete account of the models and their performances are listed in Supplementary Table 2.

**Figure 2.**
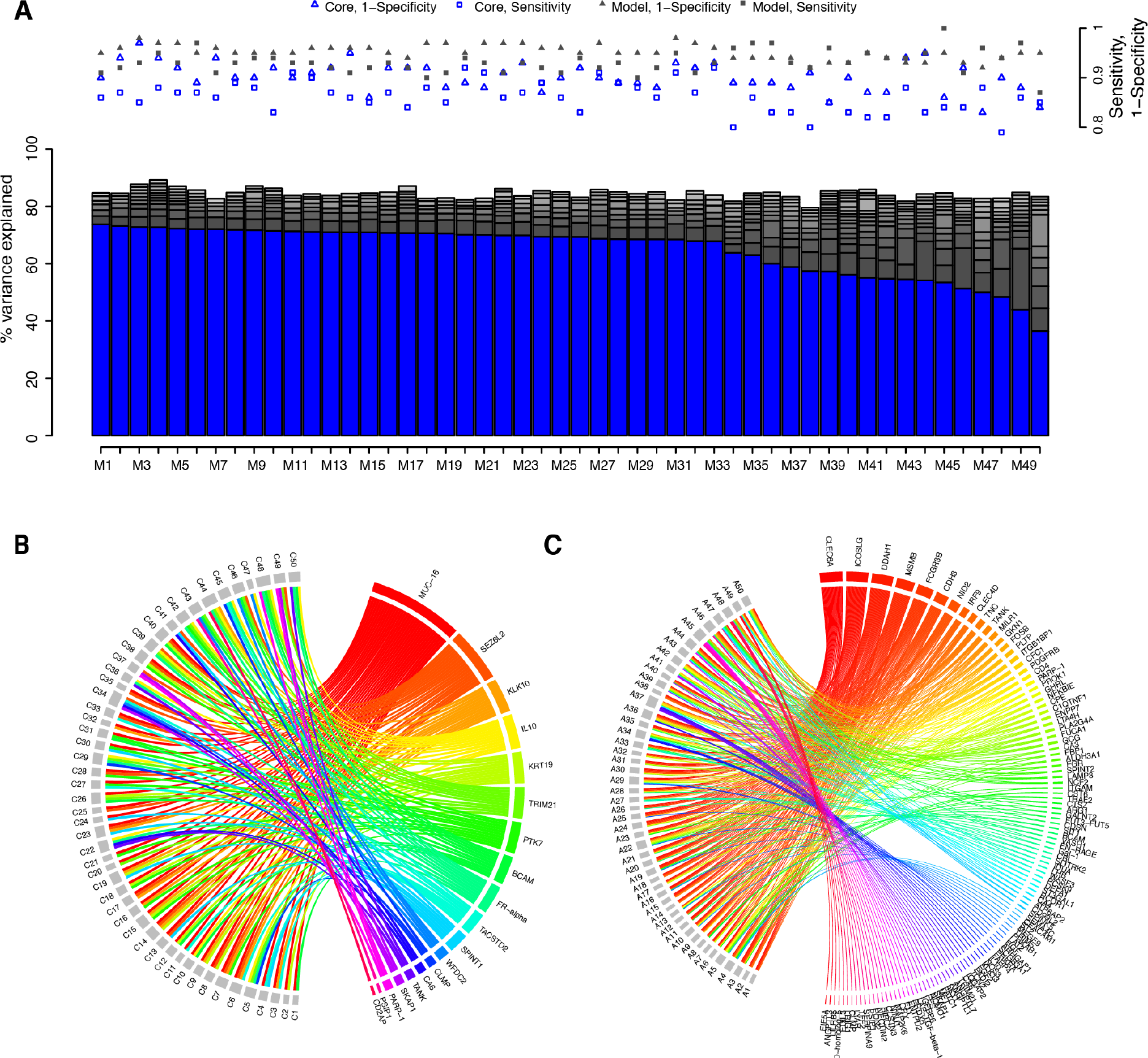
Top 50 model characteristics. (A) Variance explained in the decision (Benign tumour or Ovarian Cancer Stage III-IV by the cores (as indicated in blue) and by the additional proteins (grey) in the test set of the Discovery Data. Sensitivity and 1-Specifity of the cores (hollow markers) and the full models (filled markers) are shown (right axis) in red. **(B)** Protein inclusion into cores. Top 50 cores are indicated with C1, …, C50 and proteins are labelled with their short name. A connector represents inclusion of that protein in a core. **(C)** Same as (B) but for additional proteins (not including core-proteins). Top 50 additional protein-sets are indicated by A1, …, A50.

**Table 1:** Cohort statistics.

| Cohort | Origin<sup>1</sup> | Types | No of Women | Age, mean (SD) | CA-125<sup>2</sup> |
| --- | --- | --- | --- | --- | --- |
| Discovery | Gbg | Benign Tumours | 90 | 60.0 (16.8) | 16.8 (9.9) |
|  |  | Stage I-II | 42 | 60.7 (12.4) | 67.6 (72.0) |
|  |  | Stage III-IV | 37 | 63.8 (14.1) | 327.4 (284.5) |
| 1<sup>st</sup> Replication | Gbg | Benign Tumours | 71 | 60.2 (14.5) | NA |
|  |  | Stage I-II | 44 | 62.4 (13.7) | NA |
|  |  | Stage III-IV | 56 | 61.6 (11.3) | NA |
| 2<sup>nd</sup> Replication | UCAN | Stage I-II | 13 | 55.9 (15.0) | NA |
|  |  | Stage III-IV | 64 | 59.4 (12.0) | NA |

<sup>1</sup> UCAN: collection at Uppsala Biobank, Uppsala University, Sweden. Gbg: Gynaecology tumour biobank at Sahlgrenska University Hospital, Göteborg, Sweden <sup>2</sup> Measured at clinic [kU/L], median (median absolute deviation). NA indicates 'not available'.

**Table 2:**
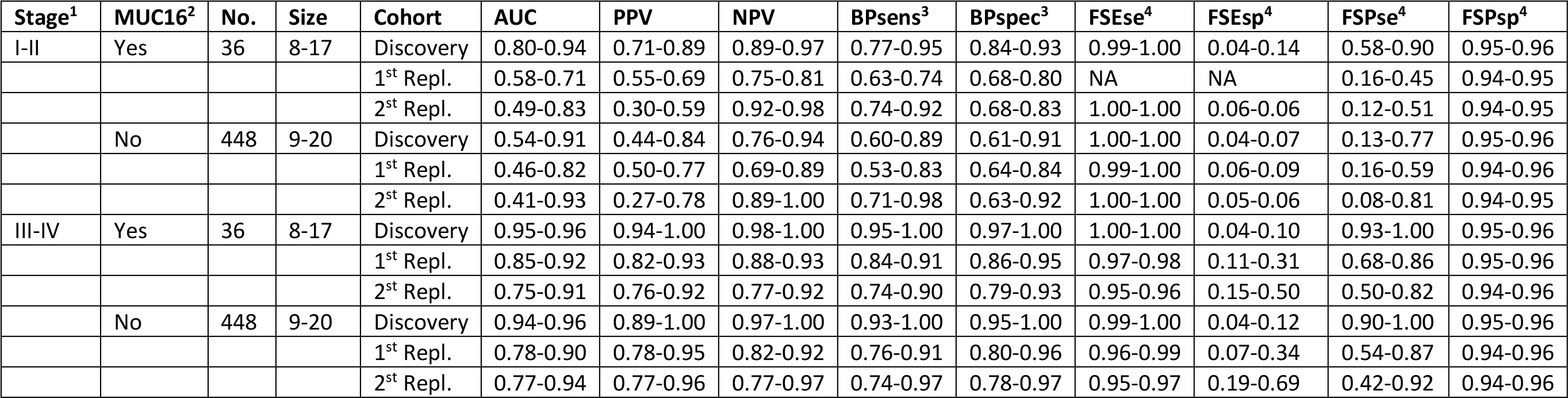
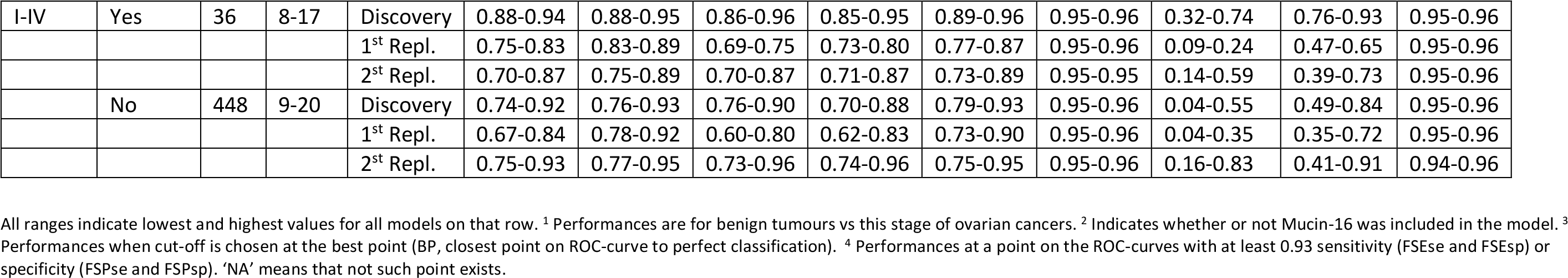
Performance ranges of all models

### Replication of model performance

The performance of each model created from the discovery data was then evaluated in two replication cohorts. As the second replication cohort lacked patients with benign tumours, the benign tumours from the first replication cohort was used in both replication cohorts. Due to the relativeness of the NPX-scale (Material and Methods) and that the data in the discovery and replication sets were generated in different laboratory analysis runs, including parts of the data that was generated using a custom-panel [Enroth et al, unpublished], the replication cohorts were split into a test and training set (50-50) and model coefficients were re-determined with the R-package ‘glmnet’^10^. The performance of the models was then estimated in the training set. This was repeated 50 times for each model and the mean and standard deviation of sensitivity, specificity, positive and negative predictive values (PPV/NPV) and AUCs were recorded. The sensitivity and specificity were calculated at three different points on the ROC curve. The ‘best point,’ defined as the closest (Euclidean distance) point to perfect classification, and by selecting a minimum sensitivity or specificity of 0.93. The performance ranges of the models are listed in Table 2. The top-ranking model all contained MUCIN-16, but overall, the average performance of models with MUCIN-16 did not display any pattern in terms of improved result relative to those without MUCIN-16. About one third of the categories showed statistically higher scores in models with MUCIN-16, about one third had lower scores and the last third did not show any significant difference in score (Wilcoxon ranked sum test, Bonferroni adjusted p-values, Supplement Table 3).

### Top-ranking model

The top-ranking of the 484 models was based on a three-protein core with MUCIN-16, TACSTD2 and SPINT1. This core was extended with 11 additional proteins (FCGR3B, TRAF2, GKN1, CST6, SEMA4C, NID2, CEACAM1, CLEC6A, MILR1, CA3 and CDH3). The distribution of abundance levels for the core proteins in the 1^st^ replication in patients with ovarian cancer stages III-IV and those with benign tumours are shown in Figure 3A. The core proteins have clearly deviating levels between the cancer cases and controls. This is further illustrated by a principle component analysis (PCA) based on the three core proteins (Figure 3B). The additional proteins are selected based on explained variance in the decision after adjustment for the variance explained by the already included proteins (Material and Methods). Therefore, some of the first six additional proteins (Figure 3C) do not differ significantly in abundance between cases and controls when examined separately but contribute to the separation when examined in combination with the previously included proteins. The separation between benign tumours and ovarian cancer stages III-IV for the top-ranked 14-protein model is shown in the PCA in Figure 3D.

**Figure 3.**
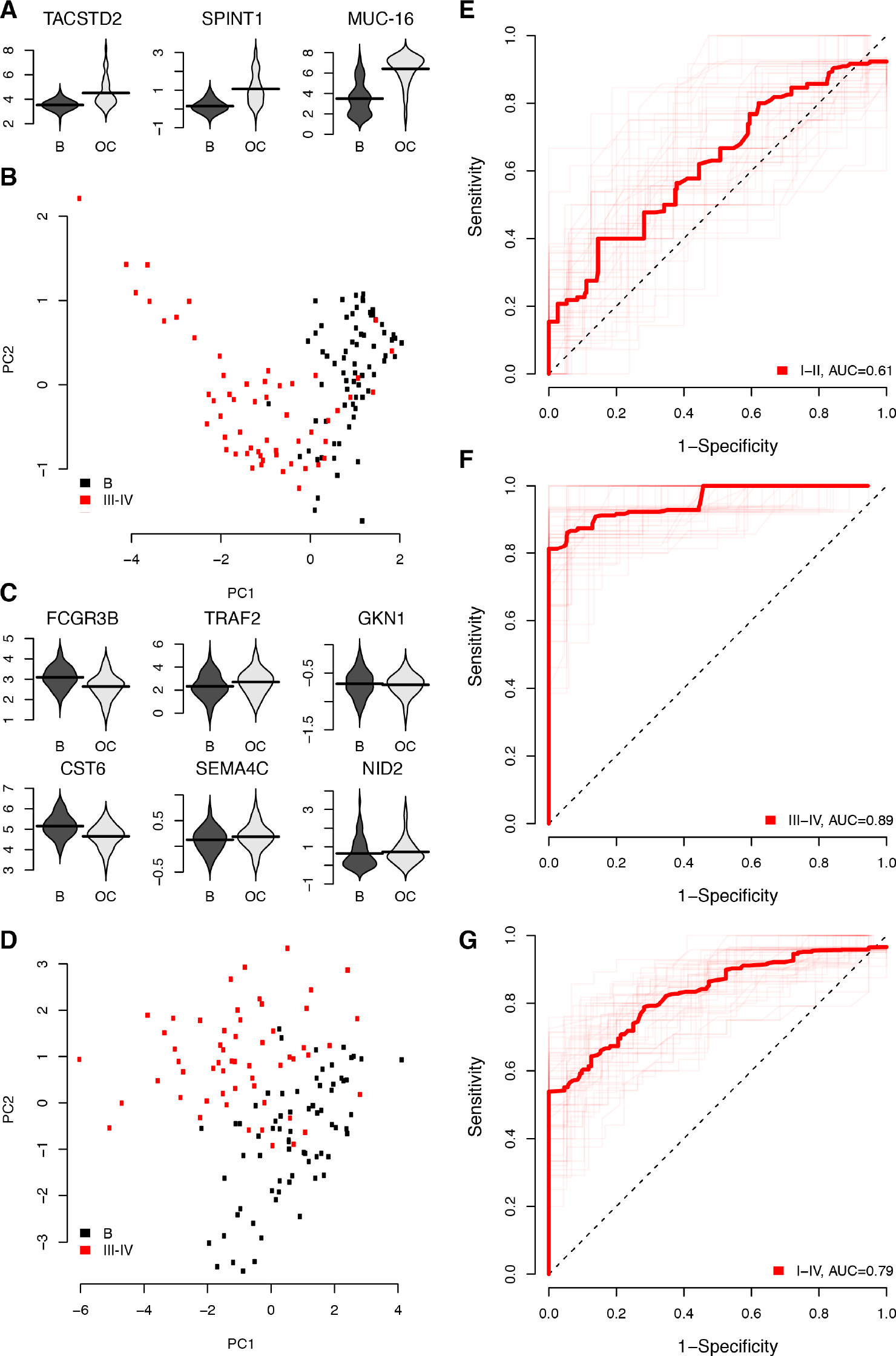
Top-ranking model performance in 1^st^ replication cohort. **(A)** Distribution of protein abundance levels in NPX for the three proteins in the core in patients with Benign tumours (indicated with a ‘B’) and Ovarian Cancer stage III-IV (indicated with ‘OC’). **(B)** PCA plot of the first two components using the proteins in the core. Figures shows Benign tumours in black and Ovarian Cancer stages III-IV in red. **(C)** As (A) but for the six first additional proteins in the model. **(D)** As (B) but for the complete model with 14 proteins. **(E-G)** Receiver Operating Characteristic (ROC) curves of the performance of the complete model in the 1^st^ replication cohort. From top to bottom, the ROC-curves represent Benign tumours vs. Ovarian cancer stages I-II, III-IV and I-IV respectively.

Receiver Operating Characteristic (ROC) curves for benign tumours versus ovarian cancer stages I-II, III-IV and I-IV are shown in Figure 3E-G. Similar illustrations for the discovery and 2^nd^ replication cohort are given as Supplementary Figure 1 and 2. For separating benign tumours from ovarian cancer stages III-IV, the top-ranked 14-protein model had an AUC=0.9, a sensitivity=0.99 and a specificity=1.00 in the test-proportion of the discovery data. In the test proportion of the 1^st^ replication data the model had an AUC=0.89, a PPV=0.93, a sensitivity=0.89 and a specificity=0.95. This should be compared to MUCIN-16 which by itself had an AUC=0.70, a PPV=0.81, a sensitivity=0.86 and a specificity=0.85 in same cohort (Figure 3F, Table 3). At a sensitivity above 0.93 in 1^st^ and 2^nd^ replication cohorts, the model achieved a specificity of 0.27 and 0.28, respectively, and at a specificity above 0.93 a sensitivity of 0.86 and 0.80. Performance measures for the discovery and replication cohorts for all the different stages investigated are listed in Table 3.

**Table 3:**
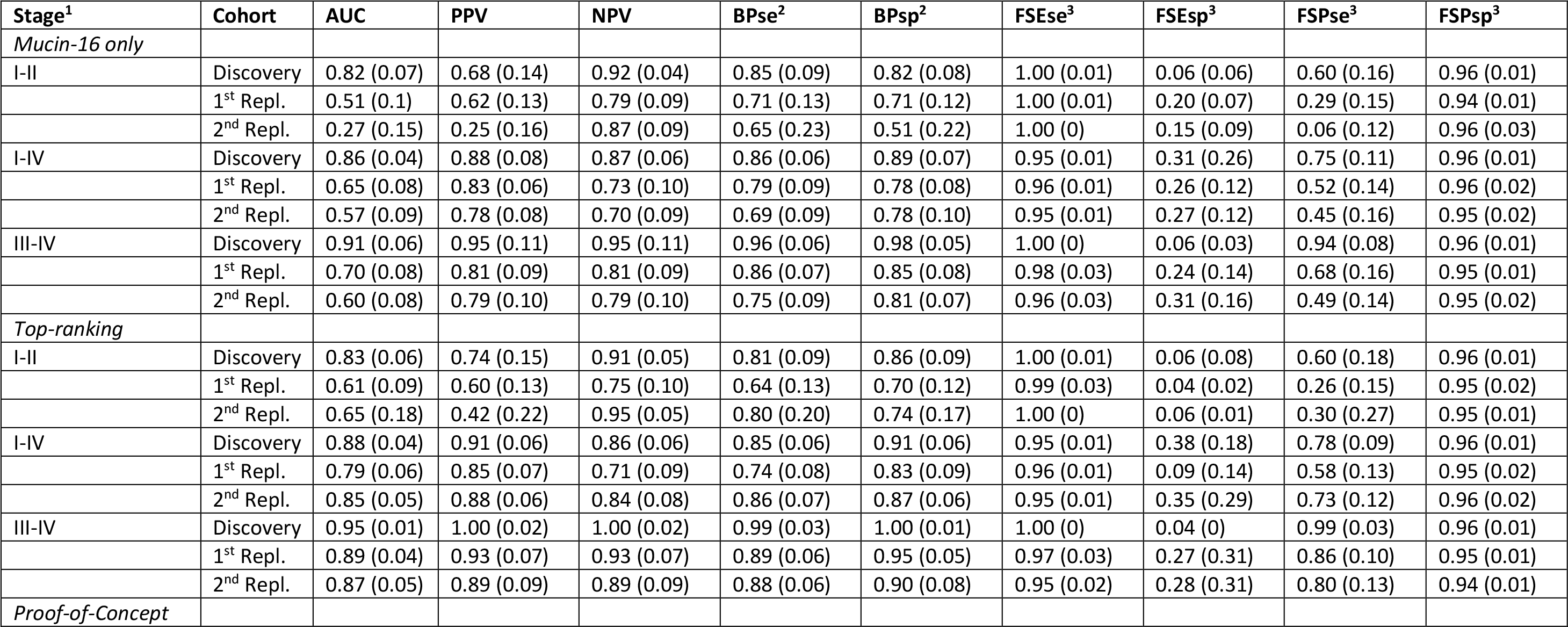
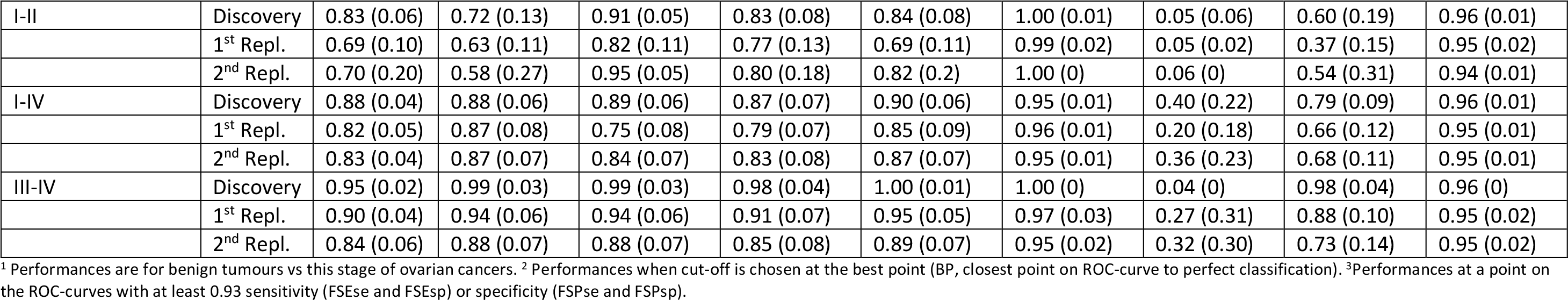
Performance of the top-ranking and the proof-of-concept model

### Proof-of-concept model for practical use

Several factors in addition to the ability to separate cases and controls may influence the choice of the proteins included in a multiplex test, such as comparison with established tests, measurable concentration range, and sensitivity of proteins to haemolysis of red blood cells causing leakage of proteins into the plasma. Taking these limitations into account, we started from the top-ranking core of the 484 models and allowed additional selection but restricted the search to proteins present in models with the highest performance in the discovery cohort. This list of possible additions was filtered by removing proteins sensitive to exposure to hemolysate^11^ and proteins that occur in much higher concentrations in human plasma than those in the selected core, and therefore would need to be diluted before assayed with PEA^11^. We then performed model selection as before based solely on the discovery data (benign tumours versus ovarian cancer stages III-IV) and identified a model consisted of 8 proteins. We finally added three proteins (WFDC2, KRT19 and FR-alpha) based on their previous association with ovarian cancer stages I-II in our modelling, or in previous literature^9,12,13^. The 11-protein panel consisted of MUCIN-16, SPINT1, TACSTD2, CLEC6A, ICOSLG, MSMB, PROK1, CDH3, WFDC2, KRT19 and FR-alpha. The performance of this 11-protein panel was then evaluated in the two replication cohorts (Table 3). In the 1^st^ replication cohort the AUC=0.90, PPV=0.94, sensitivity=0.91 and specificity=0.95 to distinguish benign tumours from ovarian cancer stage III-IV.

## Discussion

The current study was designed to identify mutually exclusive predictive biomarker signatures containing up to 20 proteins differentiating benign conditions from ovarian cancers at different stages, grades and all histological subtypes. This was done starting from a very large number of plasma proteins. These proteins were not selected based on prior association with ovarian cancer, but because of their availability in high-throughput multiplexed proteomics assays. The models were developed using a discovery cohort, and the performance of the models was then evaluated using two replication cohorts. In addition to the 484 biomarker signatures obtained using our computerized strategy, we developed one model taking into account protein-specific criteria such as abundance range and sensitivity to haemolysis. Finding combinations of predictive, robust, biomarkers is computationally intensive, and with several hundreds of proteins, exhaustive searches of combinations of up to 20 proteins is not feasible. To this end, we developed a strategy for identification of highly predictive unique signatures using hierarchical exclusion of individual proteins. By design, this lead to the discovery of many signatures that did not contain MUCIN-16. Overall, the signatures without MUCIN-16 contained a higher number of proteins than signatures with MUCIN-16, but there were no clear patterns were either group outperformed the other. Our top-ranking model achieved a sensitivity of 0.99 and specificity of 1.0 in the test proportion of the discovery data for separating benign tumours from ovarian cancer stage III-IV. A recent study by Boylan and colleagues^9^ reports perfect classification of benign tumours and late stage ovarian cancer using either MUCIN-16 or WFDC2 alone, by analysis of a single cohort with proteins measured using the same PEA technology as in our study. In our 1^st^ replication cohort, MUCIN-16 alone had an AUC of 0.70, 0.65 and 0.51 for separating benign tumours from ovarian cancer stages III-IV, I-IV, and I-II, respectively (Figure 3F-G). The difference in performance between our study and that by Boylan and colleagues^9^ could be due to geographic origin of the cohorts (USA and Sweden), biological nature of the sample (i.e. serum versus plasma), or differences in sample sizes and model evaluations. Boylan and colleagues^9^ used 21 women with benign conditions and 21 with late stage ovarian cancer, as compared to 71 and 56 in our study. Another study by Han and colleagues^8^ reported a sensitivity of 0.87 at a specificity of 1.0 for separating benign tumours from ovarian cancer stage I-IV, using the four proteins MUCIN-16, E-CAD, WFDC2, and IL-6. Our top-ranked model had a sensitivity of 0.85 and specificity of 0.91 under the same conditions. Similar to the results of these previous studies^8,9^, the performance of our models in the test-proportion of the discovery data is very good, with some models showing perfect classification. We also evaluated the selected models in two replication cohorts and found the performance similar, but somewhat lower than in the discovery set. This either implies that there are underlying differences between the cohorts, such as pre-analytical conditions, or that the models are over-trained with respect to the samples in the discovery cohort. The performance in the test-proportion of the discovery cohort should therefore be considered less certain than the results obtained in the replication cohorts. In our study, the benign tumours and the cancer samples from the 2^nd^ replication cohort differ in pre-analytical context, which could explain part of the lower performance as compared to using the 1^st^ replication cohort. This highlights the importance of understanding the context in which a biomarker test is to be used as compared to the setting used for development of the model.

Some of the proteins in the 14-protein panel and the 11-protein panel, aside from MUCIN-16 and WFDC2 (HE4), have previously been associated to ovarian cancer. TACSTD2 (Tumor-associated calcium signal transducer 2) expression has been associated with decreased survival of ovarian cancer and proposed as a prognostic factor^14^, and a biomarker for targeted therapy^15^. SPINT1 (Matriptase, HAI-2) is a type II transmembrane serine protease expressed on epithelial ovarian tumour cells. In advanced stage ovarian tumours, matriptase is expressed in the absence of HAI-1, its inhibitor, indicating that an imbalance between matriptase and HAI-1 is important in the development of ovarian disease^16^. Matriptase has also been proposed as an adjuvant therapeutic target for inhibiting ovarian cancer metastasis^17^. TRAF2 (TNF receptor-associated factor 2) regulates activation of NF-kappa-B and JNK and is involved in apoptosis. Genetic polymorphisms in this gene have been associated with high-grade serous ovarian cancer and patients with clear cell ovarian carcinoma^18^. NID2 (Nidogen-2) has been proposed as blood biomarker for ovarian cancer and is strongly correlated to CA125 levels^19^. CEACAM1 (Carcinoembryonic antigen-related cell adhesion molecule 1) is a cell-cell adhesion receptor and strongly expressed in malignant ovarian tumours^20^. Analysis of circulating tumour cell RNA have seen an increased expression of KRT19 (Keratin, type I cytoskeletal 19), but no studies of the plasma protein level have been performed^21^. FR-alpha (Folate receptor alpha, FR-alpha) is a GPI-anchored glycoprotein and serum levels has been found to be elevated in ovarian cancer patients^22,23^ and correlated to both clinical stage and histological type^24,25^. Finally, decreased expression of MSMB (Beta-microseminoprotein) has been shown to correlate with reduced survival of invasive ovarian cancer^26^. In order to study the potential of using the protein panels in diagnosis or screening, we determined their performance after tuning the models prioritizing either specificity or sensitivity. A diagnostic test for women with a transvaginal ultrasound indication of adnexal mass must possess a high sensitivity but can accept a moderate specificity. Previous investigations predicting the risk of malignancy in adnexal masses using TVU only^27^, reports sensitivities ranging from 99.7% to 89.0% with specificities of 33.7% to 84.7% for calculated risk scores of 1 to 30% and positive predictive values ranging from 44.8 to 75.4%. At a minimum sensitivity of 0.93 (actual sensitivities 0.97 and 0.95 in the 1^st^ and 2^nd^ replication cohorts) our top-ranked protein model can distinguish between women with benign tumours and ovarian cancer stage III-IV with a specificity of 0.27-0.28 and positive predictive values of 0.93-0.89. An earlier report^28^ retrospectively examined the predictive value of MUCIN-16 and WFDC2 among Swedish women that underwent surgery with suspected ovarian cancer. Out or 373 women, 58% were found to have benign tumours and 30% have ovarian cancer (15% stage I-II, 15% stage I-IV). That study reported a sensitivity of 61.9% at specificity of 75% with a positive predictive value of 31.3% for MUCIN-16 and WFDC2 combined. Thus, the performance measures of the model presented here are higher than the current clinically used biomarker combinations, but lower than the highest reported performances of clinical specialists, albeit with a higher positive predictive value. A combined use of both TVU and a biomarker test is likely to give even higher specificity. An indication of the potential for using the protein model for identification of women at risk in population screening was obtained by studying the sensitivity at high specificity. At a minimum specificity of 0.93, the top-ranked 14-protein panel has sensitivity for stages III-IV of 0.86/0.80 (1^st^ /2^nd^ replication cohort) and for stages I-IV 0.58/0.73 in distinguishing benign tumours from women with ovarian cancer (Table 3). Further studies are needed using samples collected at different time-points prior to diagnosis to evaluate the potential of using the panel in population screening. In screening, the aim is not to distinguish between benign tumours and cancer, but between healthy women and cancer, and it is likely that there will be more pronounced differences when comparing to a healthy population. In support of this notion, we have shown^29^ that the sensitivity to distinguish population controls from stage I-IV cancer was 0.62 and stage III-IV was 0.78. Future studies including age-matched population controls have to be conducted to determine the performance of the 14-protein biomarker set in population screening.

In summary, we have developed a strategy for identification of protein cores that resulted in mutually exclusive combinations of protein signatures that can separate between benign tumours and ovarian cancers. The results demonstrate the ability to achieve high performance characteristics without including MUCIN-16. We also show that broad searches for novel combinations of protein biomarkers that on their own are not necessarily good predictors is a powerful approach for finding relevant biomarkers for disease.

## Materials and Methods

### Samples

Plasma samples of women with benign and malignant ovarian tumours, either came from the UCAN collection at Uppsala Biobank, Uppsala University, Sweden or the Gynaecology tumour biobank at Sahlgrenska University Hospital, Göteborg, Sweden, as previously described [Enroth et al, unpublished] (Table 1). All tumours were examined by pathologist specialized in gynaecologic cancers for histology, grade and stage according to International Federation of Gynaecology and Obstetrics (FIGO) standards. All plasma samples were frozen and stored at −70°C. The study was approved by the Regional Ethics Committee in Uppsala (Dnr: 2016/145) and Gothenburg (Dnr: 201-15).

The discovery cohort consisted of 90 patients diagnosed with benign tumours and 79 patients with ovarian cancer stages I-IV. Samples were collected at time for primary surgery under full anesthesia but before incision. All women had at least 6 hours fasting before sample collection. The first replication cohort consisted of 71 patients diagnosed with benign tumours and 100 patients with ovarian cancer stages I-IV and were collected under the same conditions as the discovery cohort. The second replication cohort consisted of 77 patients with ovarian cancer stages I-IV. The second replication samples were collected at time of diagnosis, from awake patients, by a trained nurse.

### Protein measurements

We have previously quantified 460 proteins from the Olink Multiplex Cardiovascular II, Cardiovascular III, Inflammation, Neurology and Oncology panels in the discovery cohort using the proximity extension assay (PEA) [Enroth et al, unpublished]. Forty-two of these have also been quantified in the replication cohorts using PEA in two custom-design 21-plex panels^30^. Here, an additional 552 proteins were analysed using the Olink Multiplex Cardiometabolic, Cell Regulation, Development, Immune Response, Metabolism and Organ Damage panels and real-time PCR using the Fluidigm BioMark™ HD real-time PCR platform^31^ in the discovery and replication cohorts. A complete list of the 1012 assays corresponding to 981 unique proteins are listed in Supplementary Table 1. The samples were randomized across plates and normalized for any plate effects using the built-in inter-plate controls according to manufacturers’ recommendations. The PEA gives abundance levels in NPX (Normalized Protein eXpression) that is on log2-scale. Each assay has an experimentally determined lower limit of detection (LOD) defined as three standard deviation above noise level. Here, all assay values below LOD were replaced with the defined LOD-value. Samples and proteins that did not pass the quality control were removed. After quality control, 42 proteins from the custom panels and 551 from the additional 6 panels were kept. Assay characteristics including detection limits calculations, assay performance and validations are available from the manufacturer (www.olink.com).

### Model generation

First, the discovery set was randomly split into a training set and a test set with 50% of the samples in each, and a linear regression model was generated with the R-package ‘glmnet’^10^ with ‘alpha’ = 0.9 and optimized using 10-fold cross-validation in the training-set as implemented by the ‘cv.glmnet’-function. This was repeated 50 times with new train/test sets and a core consisting of the proteins present in at least 70% of the generated models was selected. In order to find mutually exclusive cores, the core-generating process was repeated in a recursive manner, excluding one protein at a time from the previous core from the available protein pool. For each newly generated core, the process was then repeated unless the core contained more than a specific number of proteins or had a sensitivity or specificity below a specified cut-off. For each new search, all previously excluded proteins were made unavailable to the current selection. The search was cancelled if more than 20 proteins had been excluded. The core-discovery process is outlined in Figure 1A and 1B. For each core, proteins were added creating a final model in a stepwise forward selection. First, the variance in the decision explained by the core was removed by keeping the residuals from a linear model generated with the protein values in the core as input and the decision as output. Then, the variance explained by any other available protein in the adjusted outcome was calculated and the protein explaining the most remaining variance in the decision was added to the model and the contribution of that protein to the explained variance in the decision was adjusted for. This was repeated until the best candidate protein did not explain more than 1% of remaining variance or the total number of proteins in the model exceeded 20 proteins (Figure 1C).

## Author contribution

UG is study PI. SE, KSU and UG designed the study. KSU, KS, MO and MLy contributed patient material. MLi and EA generated protein data and performed quality control. SE, MB developed analysis tools and performed computational analyses. SE, JB, MB, KSU and UG interpreted data. SE and UG drafted the manuscript. All authors contributed in the writing of the final version of the manuscript.

## Acknowledgements

The study was funded by the Swedish Cancer Foundation, The Swedish Foundation for Strategic Research (SSF), the Swedish Research Council (VR), VINNOVA (SWELIFE) and Olink Proteomics.

